# SIRP: A Self-Supervised Fluorescence Denoising method with Implicit Representation Priors

**DOI:** 10.1101/2025.11.25.690616

**Authors:** Yuze Li, Jun Zhu, Pengcheng Xu, Min Guo, Yue Li, Huafeng Liu

## Abstract

Fluorescence microscopy plays a critical role in live-cell imaging, yet the captured images often suffer from significant noise due to limited excitation intensity and constrained exposure times. Deep learning has emerged as a powerful solution for image denoising, especially the self-supervised methods can achieve good restoration performance without requiring clean target data. However, their performance in preserving fine image details remains challenging under low signal-to-noise ratio (SNR) conditions. In this work, we propose a novel self-supervised denoising framework named Self-Supervised Implicit Representation Prior Image Restoration Network (SIRP). It is based on the neighbor-sampling strategy and leverages image representation priors to guide the network training procedure. The neighbor-sampling strategy takes advantage of the inherent statistical consistency within local image regions to denoise. The image representation priors are learned through the Implicit Neural Representations (INR) that map spatial coordinates directly to their corresponding pixel intensities to suppress noise and effectively preserve details. Results of SIRP on simulated data and experimental-acquired data showed superior denoising performance and detail-preserving ability compared to filter-based approaches and self-supervised approaches, highlighting the potential of INR-guided architectures for fluorescence microscopy image restoration.

## 1. Introduction

Live cell imaging has become an indispensable tool in modern biological research. It enables the direct observation of dynamic cellular processes in their native environments, thereby enhancing the understanding of fundamental biological mechanisms. Among the various available techniques, fluorescence microscopy [1] has emerged as one of the most widely used tools for live cell imaging, due to its high specificity and high contrast [2].

However, fluorescence microscopy inherently involves a trade-off between spatial resolution, temporal resolution, and phototoxicity [3–6]. Achieving high-quality images often requires strong excitation and long exposure times, but these conditions exacerbate photobleaching and induce phototoxicity that can compromise cell viability. To minimize such adverse effects, live imaging experiments are typically performed under reduced illumination and shortened exposure times [7].

As a result, fluorescence microscopy under such low-light-dosage conditions inevitably suffers from a reduced signal-to-noise ratio (SNR), where noise overwhelms the acquired signal and compromises image quality. The degradation in SNR not only obscures fine structural details but also hinders downstream analysis tasks such as segmentation, tracking, and quantitative measurement [8]. Therefore, effective denoising strategies are crucial preprocessing techniques for improving image quality in low-light live-cell imaging.

Traditional image denoising techniques—such as Gaussian filtering [9], median filtering, bilateral filtering [10], and non-local means (NLM) [11]—have long been valued for their simplicity, efficiency, and ability to preserve key image structures. Gaussian and median filters reduce high-frequency and impulsive noise, while bilateral filtering adaptively smooths intensity variations without blurring edges [9, 10]. NLM further exploits self-similar patterns across the image to achieve strong denoising performance with minimal structural distortion [11], and has been successfully applied in applications such as live-cell imaging in Drosophila [12] and zebrafish larvae brain imaging [13]. However, despite their practicality, these methods often struggle with the complex noise characteristics and extremely low SNR typical of fluorescence microscopy: Gaussian, median, and bilateral filters tend to oversmooth fine subcellular features, and NLM may still lose weak signals without careful adaptation [12, 14].

With the rise of deep learning, data-driven methods have received significant attention for image denoising. Deep learning–based denoising approaches can generally be divided into supervised and unsupervised categories. Supervised methods rely on paired noisy-clean datasets for training, enabling precise noise removal [15, 16]. However, the acquisition of clean ground-truths is costly and sometimes impractical, due to sample non-repeatability, photobleaching, and phototoxicity constraints [5]. Although synthetic datasets may alleviate this limitation, they often suffer from domain gaps that fail to generalize well to real-acquired data [17], restricting the applicability of supervised models.

To overcome these challenges, self-supervised and unsupervised frameworks have been proposed. These strategies can be broadly grouped into several classes. Neighbor/pair-based approaches (Noise2Noise [18], Neighbor2Neighbor [19], Noise2SR [20]) exploit multiple noisy realizations or local pixel sampling, but their performance hinges on noise independence or repeated acquisitions, which may not hold in structured or dynamic fluorescence imaging. Blind-spot approaches (Noise2Void [17], Noise2Self [21], DeepSeMi [22]) mask pixels during training to avoid trivial identity mappings; however, masking discards inevitably observed information at those locations, potentially causing loss of fine or high-contrast structures. Other methods adopt sub-sampling or architectural modifications such as asymmetric or singular convolutions [22, 23], but these approaches may still compromise high-frequency structures through information loss or aliasing. As a result, these strategies struggle to preserve high-frequency or fine-grained image details in low-SNR scenarios, where signal information is intrinsically scarce.

Leveraging image prior knowledge has shown potential to mitigate information loss. For example, approaches like Deep Image Prior (DIP) [24, 25] can provide meaningful structural constraints and compensate for missing data even without large annotated datasets. Therefore, designing a method that effectively embeds image knowledge into an end-to-end, self-supervised denoising framework holds positive promise for improving denoising quality under low SNR conditions.

Building on this insight, we propose the Self-Supervised Implicit Representation Prior Image Restoration Network (SIRP), a framework that utilizes implicit neural representations (INRs) [26] to provide prior information for neighbor-sampling-based denoising network training. INRs map continuous spatial coordinates to signal values rather than operating on discrete pixel grids, which gives them an inherent bias toward low-frequency and smoothly varying structures. During training, the INR image representation module branch encodes image structure and captures the underlying effective signal components to provide better detail preservation. Once training is complete, the network operates independently of the image representation module and achieves an efficient denoising process. By integrating the representational strength of INRs with the statistical consistency leveraged by neighbor-sampling, SIRP effectively suppresses high-frequency noise while maintaining morphological fidelity. Results of both simulated and experimentally-acquired data demonstrate that SIRP consistently improves image quality across a wide range of signal-to-noise ratios. This approach offers a practical and powerful solution for fluorescence microscopy denoising and is particularly beneficial for live-cell imaging.

## 2. Methods

### 2.1 Overview

The proposed Self-Supervised Implicit Representation Prior Image Restoration Network (SIRP) is designed to be trained in a two-stage process. It consists of two core modules: an Image Representation Module that extracts a clean, structure-preserving prior from a single noisy input using Implicit Neural Representation (INR), and an end-to-end denoising network that leverages this prior and neighborhood information to perform effective noise removal. The overall scheme of the proposed framework is illustrated in Fig. 1, including the training process, the inference process, and the details of the image representation module.

**Fig. 1.**
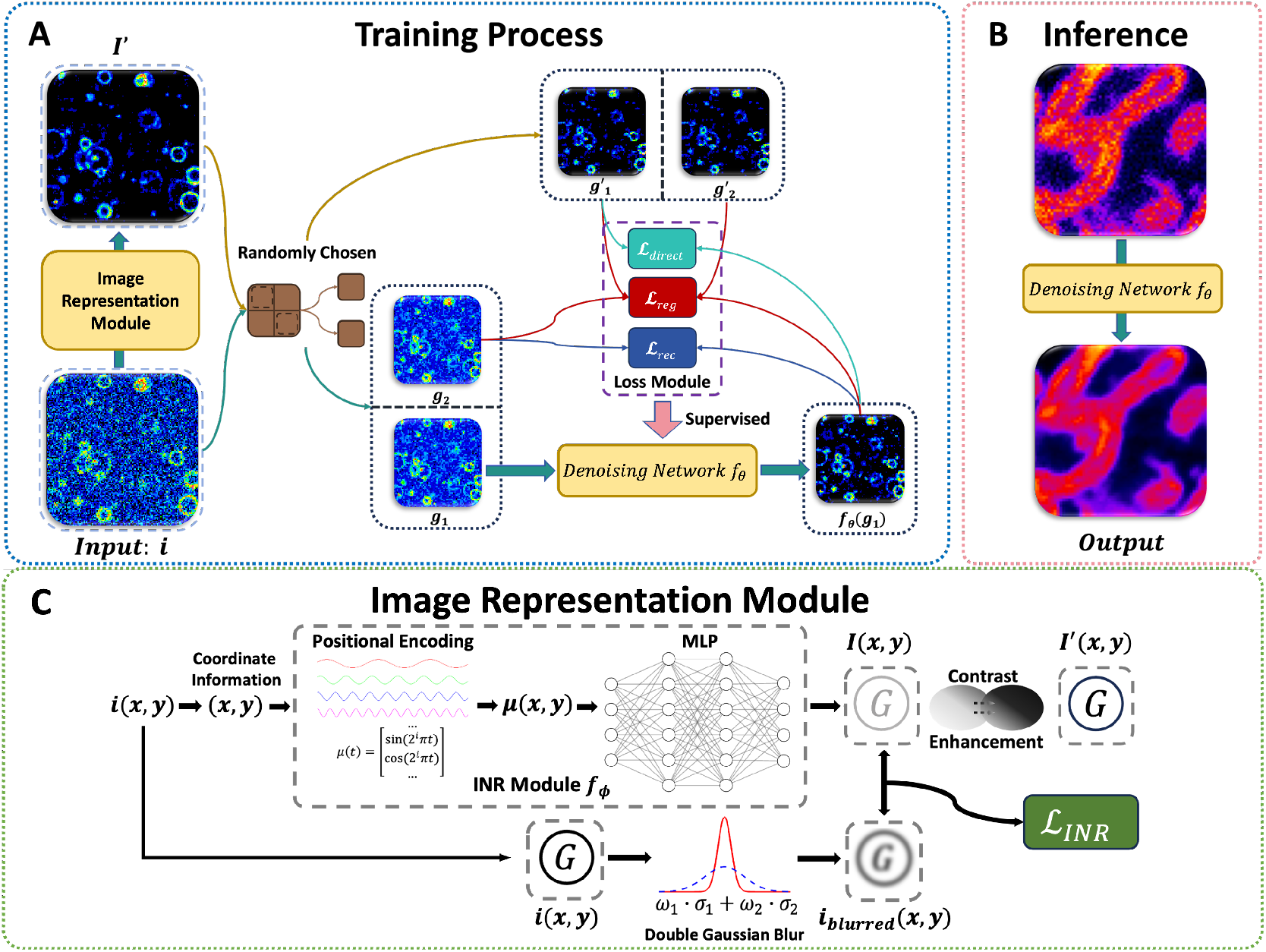
The implementation of SIRP. (A) Overview of the training process. The input image *i* is first encoded by the image representation module to obtain an image-prior-term *I*′ (*i*). Both *i* and *I*′ (*i*) are then randomly downsampled within a 2 × 2 window to generate two sets of sub-images, i.e. *g*_1_,*g*_2_ for *i* and 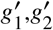 for *I*′ (*i*). These sub-images are used within the loss module to supervise the training of the denoising network. Through this iterative training process, the denoising network is progressively optimized and can subsequently be applied for practical fluorescence image restoration tasks. (B) The inference stage of the proposed framework. Once training is completed, new noisy images can be fed into the network and get the restored output. The image representation module is not used during the inference period. The example shown here is an image of mitochondria. (C) The coordinate information of each pixel location (*x, y)* from the input image (*i (x, y)* is first processed by the INR module *f*_*ϕ*_. The coordinates are mapped into a high-dimensional positional encoding (*µ (x, y)*, which is subsequently fed into an MLP to predict the corresponding intensity value (*I (x, y)*. In parallel, the input image (*i x, y)* undergoes a double Gaussian blur to smooth local variations, producing the blurred reference image *i*_blurred_ (*x, y)*. The blurring degree is controlled by the Gaussian standard deviations *σ*_1_ and *σ*_2_, together with their mixing weights *ω*_1_ and *ω*_2_. The loss function ℒ_INR_ is computed by the *L*_2_ distance between *i*_blurred_ (*x, y*) and the predicted output *I* (*x, y*), while additional *L*_1_ and *L*_2_ penalties on the output further suppress high-frequency fluctuations. Finally, the predicted image *I* (*x, y*) is refined using an exponential contrast adjustment, generating the final representation *I*′ (*x, y*).

The key innovation of the proposed method lies in the integration of image knowledge into an end-to-end, self-supervised denoising framework. The image representation module leverages the unpredictability of noise and generates a valid prior featuring a clean and consistent background, offering effective global structural guidance for the pixel-level denoising network. This design leads to a robust and consistent image restoration under low signal-to-noise conditions.

### 2.2 Image Representation Module

To generate a structure-preserving prior from a single noisy image, we employ an Implicit Neural Representation (INR), which is implemented as a multi-layer perceptron (MLP) *f*_*ϕ*_, to learn a continuous mapping from spatial coordinates to image intensities. Owing to the inherently difficult-to-learn nature of noise and the inherently structured, smooth characteristics of the underlying signal, this coordinate-based formulation enables the model to represent fine structural details while naturally suppressing pixel-wise noise.

To capture both global smoothness and fine local variations, the 2D coordinates (*x, y*) are first mapped into a high-dimensional representation using Fourier positional encoding [27]:

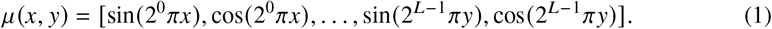

where *L* denotes the number of frequency bands, which controls the highest encoded frequency, a larger *L* captures finer spatial details, whereas a smaller *L* yields smoother representations.

This encoding allows *f*_*ϕ*_ to model multi-scale spatial frequencies effectively, enabling it to represent continuous image structures with subpixel precision. The encoded coordinates are then fed into a fully connected network with ReLU activations to generate the reconstructed intensity:

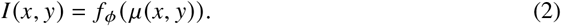

At the same time, given a noisy image (*i x, y*), we apply a double Gaussian blur to obtain a smoothed version *i*_blurred_ (*x, y*). Using two Gaussian kernels with different standard deviations allows us to balance structure preservation and background consistency: the smaller *σ* preserves essential structural details, while the larger *σ* provides stronger background smoothing. Their weighted combination reduces high-frequency noise without excessively blurring important features. The blurred image is computed as

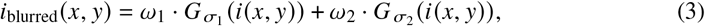

where *σ* _1_, *σ* _2_, *ω* _1_, and *ω* _2_ are hyperparameters, chosen to balance the trade-off between avoiding excessive smoothing and maintaining sufficient noise-suppression capability. These values are empirically chosen parameters in our method.

The INR parameters *ϕ* are optimized to fit the blurred image *i*_blurred_ through an *L*_2_ reconstruction loss and two output regularization terms.

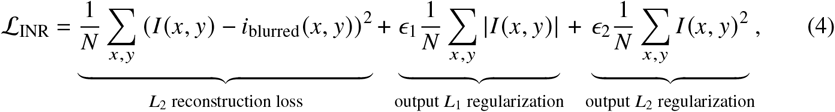

where ϵ _1_ and ϵ _2_ are tunable hyperparameters controlling the strength of the output *L*_1_ and *L*_2_ regularization, respectively.

The reconstruction term enforces pixel fidelity to the smoothed image, while the two output regularization terms provide complementary constraints [28]. The *L*_1_ regularization encourages sparsity in the reconstruction, effectively reducing unstable noise. The *L*_2_ regularization promotes global smoothness, ensuring structural stability. This joint regularization enables the model to capture the true underlying signal rather than noise fluctuations.

Although the INR can suppress the noise, the result may still appear low-contrast under low-SNR conditions. This is because the background noise in such scenarios compresses the dynamic range of the measurements, making it difficult for INR to fully restore the original contrast even after denoising. Based on the need to enhance visual and structural clarity, we apply an exponential contrast adjustment:

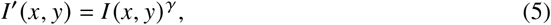

where *γ* is a tunable parameter controlling contrast intensity, a larger one increases contrast and brightens weak structures, while a smaller one reduces contrast and makes details less visible. This simple nonlinear transformation amplifies structural differences in dim regions, yielding a high-quality prior *I*′ (*x, y*) that serves as reliable supervision for the subsequent self-supervised denoising stage.

### 2.3 Self-supervised Denoising Network

The second stage of SIRP focuses on training a denoising network in a self-supervised manner, which relies solely on the intrinsic redundancy of natural images and is guided by the structural prior information (*I*′ (*x, y*) generated from the INR module. This design enables a robust training process even without paired clean-noisy datasets.

Before training, it is necessary to build a self-supervised dataset. Both the original noisy image *i* and the INR-enhanced image *I*′ are partitioned into non-overlapping 2× 2 pixel blocks. Randomly selecting one pixel in each image block for downsampling and repeating twice can create two paired downsampled views:

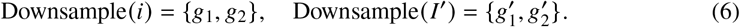

This random sub-sampling ensures that *g*_1_ and *g*_2_ contain nearly identical structures but independent noise realizations. It is similar for 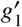 and 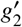. Such data construction mimics the statistical independence assumption between neighboring pixels, which is crucial for self-supervised denoising methods [17, 19, 21].

The denoising network *f*_*θ*_ aims to reconstruct underlying noise-free results by leveraging the similarity between different sub-images. Most convolutional networks can be used for this purpose. Here, we adopt a U-Net [16] architecture due to its ability to capture multi-scale features and preserve spatial resolution, which has shown strong performance in image restoration and denoising tasks. The network is optimized using three complementary loss terms, each addressing a different aspect of self-supervised learning or structural preservation.

First, we used a reconstruction loss to enforce that the prediction from one noisy image matches the other:

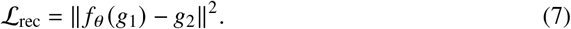

This term drives the network to recover the shared signal content between the two noisy inputs while averaging out uncorrelated noise. However, since both inputs are noisy, this loss alone may lead to residual artifacts or over-smoothing.

To address the potential instability caused by relying solely on reconstruction losses between two noisy views, which may make it difficult for the network to distinguish detailed structures from random noise fluctuations. We introduce a transformation-consistency regularization term that incorporates structural guidance from the INR prior. The key idea is that the structural relationship (i.e., the transformation) between the two sub-images derived from the “clean” INR prior provides a reliable reference for how the underlying image content should vary across different sub-images. By enforcing the network’s output to match this consistent transformation, specifically, by requiring the difference between its predictions and targets to align with the difference observed between INR-derived sub-images, we derive the loss:

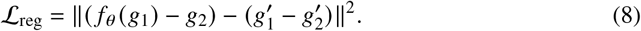

This regularization ensures that the model learns stable and smooth structural changes guided by the prior, rather than overfitting to irregular noise patterns, thereby providing a more reliable optimization trajectory.

Finally, we introduce a direct alignment objective that encourages the network output to match the INR prior at the pixel level:

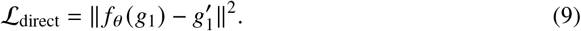

This constraint promotes spatial coherence and preserves fine structural details, leading to improved stability and cleaner background reconstructions, especially in low-SNR regions.

The overall training objective combines the above terms in a weighted form:

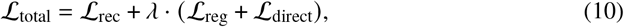

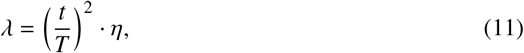

where *t* denotes the current epoch, *T* is the total number of epochs, *η* is a fixed scaling coefficient, and *λ* is a dynamically increasing weight that gradually strengthens the contribution of the INR-guided regularization as training progresses, allowing the network to first learn the basic self-supervised reconstruction before leveraging prior knowledge for fine refinement. This scheduling strategy stabilizes early optimization and leads to more reliable convergence. This hybrid loss design allows the network to benefit from both pixel-level self-consistency and structural priors, resulting in enhanced denoising performance without access to clean labels.

Overall, this two-stage training strategy—first generating reliable structural priors via INR, then guiding a self-supervised denoising network through a composite loss—enables SIRP to achieve high-quality restoration.

### 2.4 Implementation Details

All raw images are normalized individually using percentile normalization:

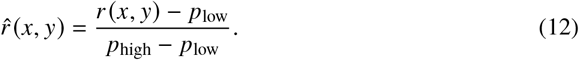

where *p*_low_ and *p*_high_ denote the 0.2-th and 100-th percentile intensity values of the image, respectively. This transformation effectively suppresses background noise while preserving the dynamic range of the fluorescence signals.

For the Image Representation Module, in all experiments, the number of high- and low-frequency components of positional encoding is fixed at 8. The INR is trained as a fully connected MLP with four hidden layers of 32 neurons each, using ReLU activations. The output layer uses a linear activation to produce continuous values. And the parameters we used in the double Gaussian blur are *σ*_1_ = 1.0, *σ*_2_ = 3.0, *ω*_1_ = 0.7, and *ω*_2_ = 0.3, respectively.

During training, each raw image is processed by a separate INR module, and the corresponding prior information is subsequently stored. The INR is trained by the Adam optimizer with a fixed learning rate of 1× 10^−3^ and a step-decay schedule (*γ* _*decay*_ = 0.95 every 100 steps). Each image is optimized for 2,000 iterations. The loss combines a mean-squared reconstruction term with *L*_1_ and *L*_2_ penalties weighted by ϵ_1_ = 0.2 and ϵ_2_ = 0.4, respectively. And the value of *η* we applied is 1.0. After convergence, an exponential contrast enhancement (*I*′ (*x, y* = *I (x, y*) ^*γ*^ with *γ* = 1.5 is applied.

The network adopts a U-Net architecture comprising six encoding stages followed by a bottleneck layer. Each encoding stage consists of one or two convolutional layers with 48 feature channels, using 3× 3 kernels, stride 1, and padding 1, and all convolutions are followed by a LeakyReLU activation with a negative slope of 0.2. Downsampling is performed using 2 × 2 max pooling, which reduces the spatial dimensions by a factor of 2 at each stage while keeping the number of feature channels constant. In the decoding path, feature maps are upsampled using 2 × 2 transposed convolutions and concatenated with the corresponding skip connections from the encoder, effectively doubling the number of channels after each concatenation. Finally, the output is generated through a series of 1 × 1 convolutions to map the 96-channel feature maps to the single-channel output.

The above parameters are chosen based on experimental validation.

## 3. Results

### 3.1 SIRP Provides Better Restoration Than Image Representation and Neighbor-sampling Method

We first evaluated our method on synthetic images. The no-noise ground-truths (GT) were generated with various geometric patterns such as circles, rings, and lines. Then they were corrupted with a mixture of Gaussian and Poisson noise to yield the raw image with SNR below 20 dB. The raw images exhibit substantial degradations: the background contrast was significantly reduced, and many structural features were partially or completely destroyed.

Since SIRP represents an integration of an image representation prior method and an end-to-end neighborhood sampling denoising strategy, we conducted a comparative study to evaluate the advantage of this combined approach over either individual strategy alone. The image representation prior (denoted as INR prior) was derived from an INR module that shared the same parameters as that of SIRP. For the neighborhood sampling strategy, we adopt the framework of Neighbor2Neighbor (Nei2Nei), which employs the same denoising architecture as SIRP but utilizes a different loss function. The performance comparison is shown in Fig. 2. Compared with the raw image, all strategies reduced the noise and provided visual quality improvements, but SIRP achieves superior restoration over using a single strategy (Fig. 2A-B). Specifically, the INR prior provides a clean background, but its details are over-smoothed and sometimes contain undesirable dark pixels (Fig. 2B). In contrast, Neighbor2Neighbor provides sharper details, but it also contains distorted backgrounds and signals, such as the broken circle boundaries in magnified views in Fig. 2B. By combining the strengths of both strategies, SIRP preserves fine structural details, maintains high-frequency components, and produces cleaner backgrounds with higher visual contrast. By leveraging an image representation prior, SIRP is guided toward more accurate structural reconstruction throughout the training process. The quantitative analysis with Peak signal-to-noise ratio (PSNR) provides a consistent conclusion with the visual comparisons (Fig. 2C). SIRP provided obvious PSNR improvements over Neighbor2Neighbor and INR prior, showing increases of more than 3 dB (31.2 dB for SIRP vs 26.0 dB for INR Prior and 27.9 dB for Nei2Nei).

**Fig. 2.**
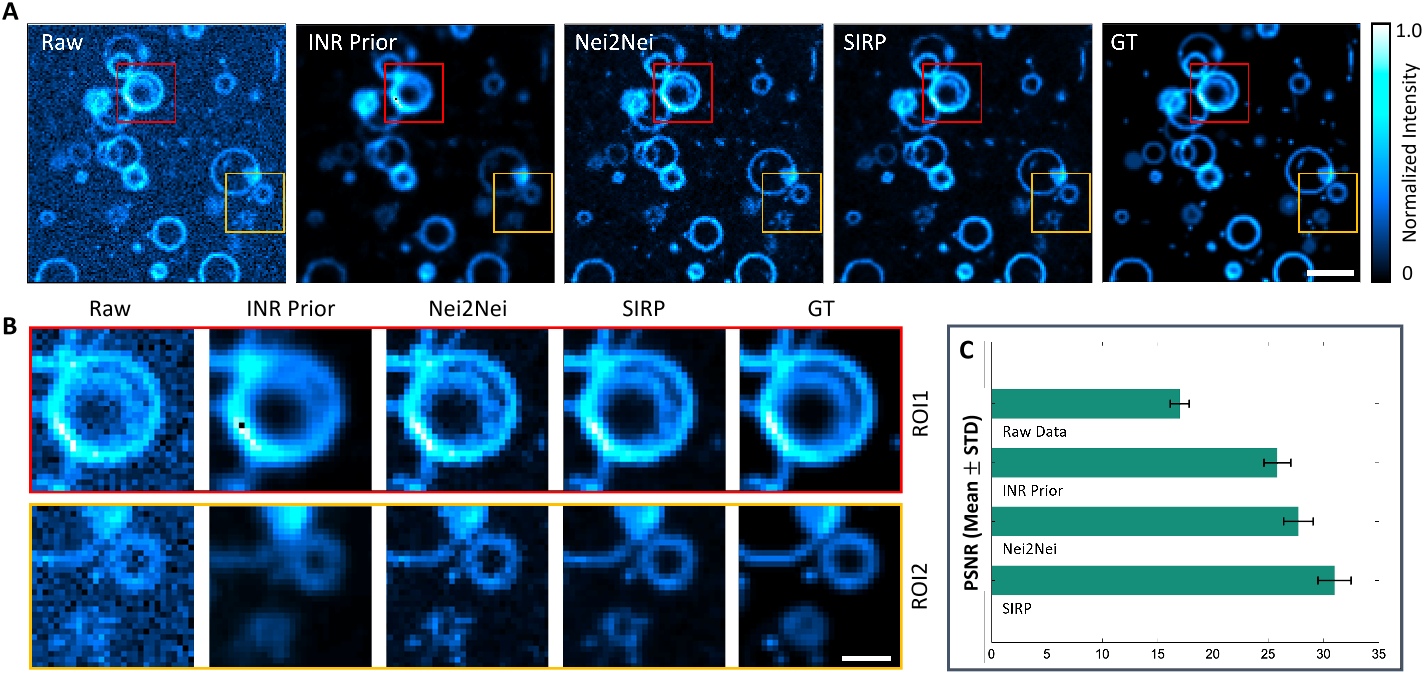
(A) Visual comparison of SIRP with INR Prior and typical neighbor-sampling-based method (Neighbor2Neighbor) on synthetic images. From left to right: Raw, INR Prior result, Neighbor2Neighbor (Nei2Nei) result, SIRP result, and ground truth (GT). SIRP produces reconstructions with a cleaner background and structural details, which are closer to GT (scale bars: 25 pixels). (B) Two regions of interest (ROIs) highlighted by red and yellow rectangles in (A), illustrating local restoration details (scale bars: 10 pixels). (C) Quantitative comparison between SIRP and other methods with PSNR (mean ± standard deviation, n=20 for analysis). SIRP provides the highest PSNR value.

### 3.2 SIRP Outperforms Traditional and Self-Supervised Denoising Methods Across Different Noise Levels

To further validate the effectiveness of the proposed SIRP framework, we compared its performance against a broad set of conventional methods and typical self-supervised algorithms on synthetic images, where different SNR levels were generated by applying different levels of Poisson noise and tuning the variance of added Gaussian noise. Conventional methods include traditional filtering (average, Gaussian, median) and a nonlocal-similarity-based method (BM3D), and the self-supervised denoisings include Noise2Void (N2V) and Neighbor2Neighbor(Nei2Nei). The comparison results are shown in Fig. 3. All methods are able to reduce noise to some extent, but their behaviors differ markedly (Fig. 3A). Traditional filtering methods show limited contrast. Median filtering enhances bright structures but leaves the background disturbed, whereas Average and Gaussian filtering oversmooth the image and blur fine details. BM3D improves structural resolution compared with these filtering approaches, yet it still fails to suppress the background. Learning-based methods achieve better performance, but N2V produces relatively blurry results with checkerboard artifacts, and Nei2Nei introduces noticeable detail distortions. SIRP produces reconstructions that are closest to the ground truth, with cleaner backgrounds and sharper boundaries than all competing methods.

**Fig. 3.**
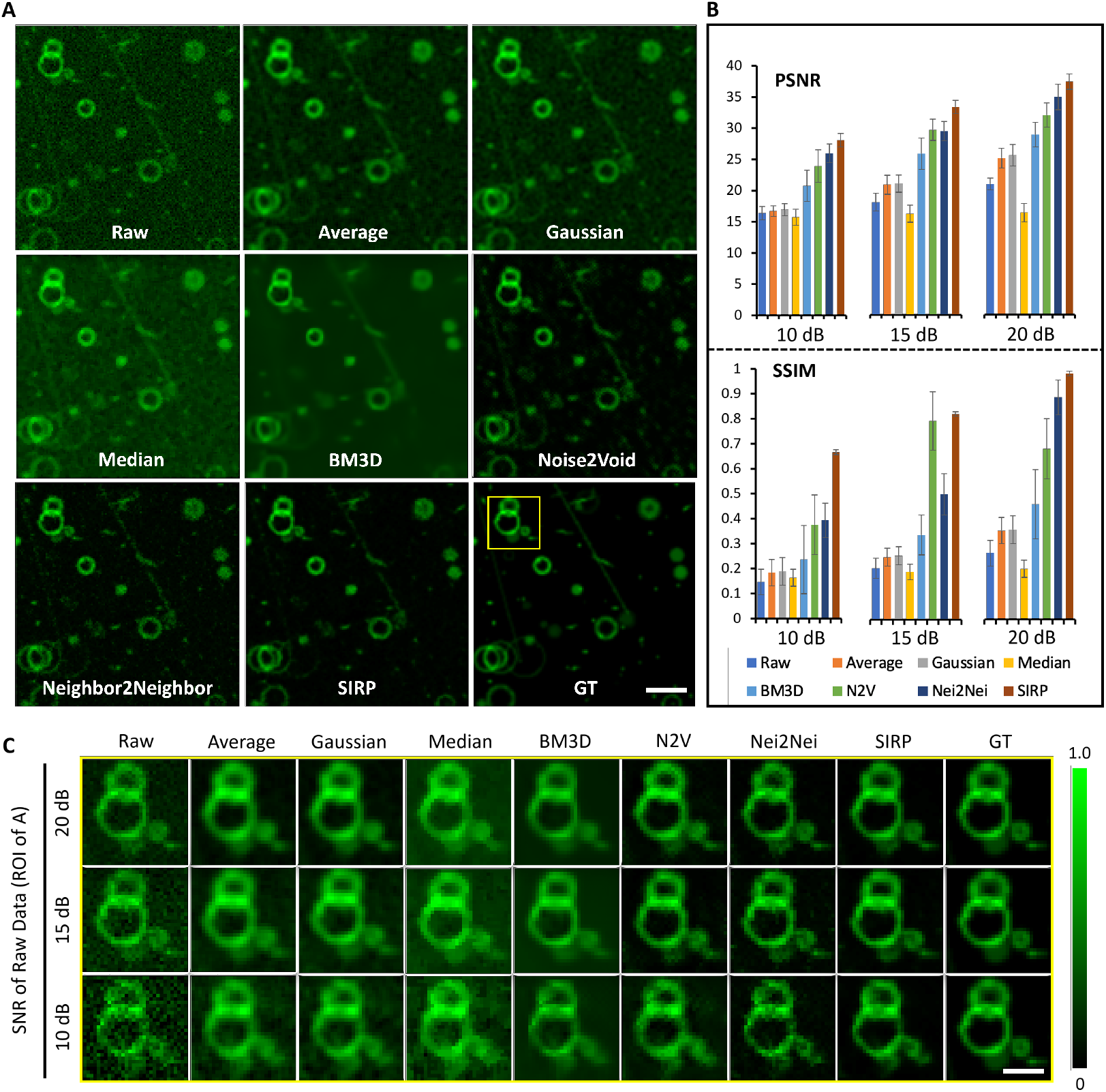
Visual and quantitative comparison of various denoising methods on simulated data under different noise levels. (A) Restoration results for Raw with SNR ∼ 15dB, Average filtering, Gaussian filtering, Median filtering, BM3D, Noise2Void (N2V), Neighbor2Neighbor (Nei2Nei), the proposed SIRP, and the ground truth (GT). The scale bar represents 25 pixels. SIRP provides good background suppression, introduces fewer structural artifacts, and achieves a resolution closer to the GT. (B) Quantitative evaluation of the seven methods across three noise levels (SNR = 10, 15, 20 dB) in terms of PSNR and SSIM. For each noise level, we compute the mean and standard deviation over 20 test images. In all cases, SIRP provides the best restoration. (C) Magnified views of ROI selected by the yellow rectangular region in (A), showing local restoration details for the seven methods under different SNR levels. Scale bar:10 pixels.

We further analyze the anti-noise stability of different methods with PSNR and structural similarity index (SSIM) metrics in Fig. 3B. Regardless of the noise level, SIRP consistently achieves high PSNR and SSIM with minimal variance, demonstrating strong robustness to noise. Importantly, although the reconstruction quality of SIRP degrades as the SNR of the raw image decreases, it delivers the best restoration results across all noise levels.

The visual comparison among different methods under different noise levels is illustrated using the magnified ROIs in Fig. 3C, where SIRP preserves local features even under severe noise like an SNR of 10dB, while other methods either oversmooth the structures or retain significant background artifacts. Collectively, these results demonstrate that SIRP provides more stable performances than other conventional and self-supervised denoising approaches across a wide range of SNR conditions.

### 3.3 SIRP Provides Robust Denoising Performance on Diverse Fluorescence Microscopy Datasets

Next, we used different real-acquired biological structures to further validate the denoising performance of SIRP, including ER, microtubules, membrane, and nucleus. We compared SIRP with Noise2Void, and the results are shown in Fig. 4. The ER and Microtubules datasets were acquired using structured illumination microscopy (SIM) [29], with low-SNR (raw) images corresponding to single-angle, low-photon-count acquisitions and high-SNR ground truth (GT) images generated by combining multiple angles or accumulating photons. For line-like structures such as ER and microtubules, SIRP preserves sharper details. Fourier spectrum analysis indicates that SIRP provides a similar high-frequency distribution as the GT and enables subtle structure reconstruction. The ‘blind-spot’ strategy used by N2V leads to inherent information loss, which consequently limits its ability to restore high-frequency details compared to SIRP (Fig. 4A). The line-profile comparisons further supports these observations, the line structures of microtubules show a narrower distribution in SIRP results, whereas Noise2Void provided blurred shapes and introduced residual background noise that disturb the line profile shape.

**Fig. 4.**
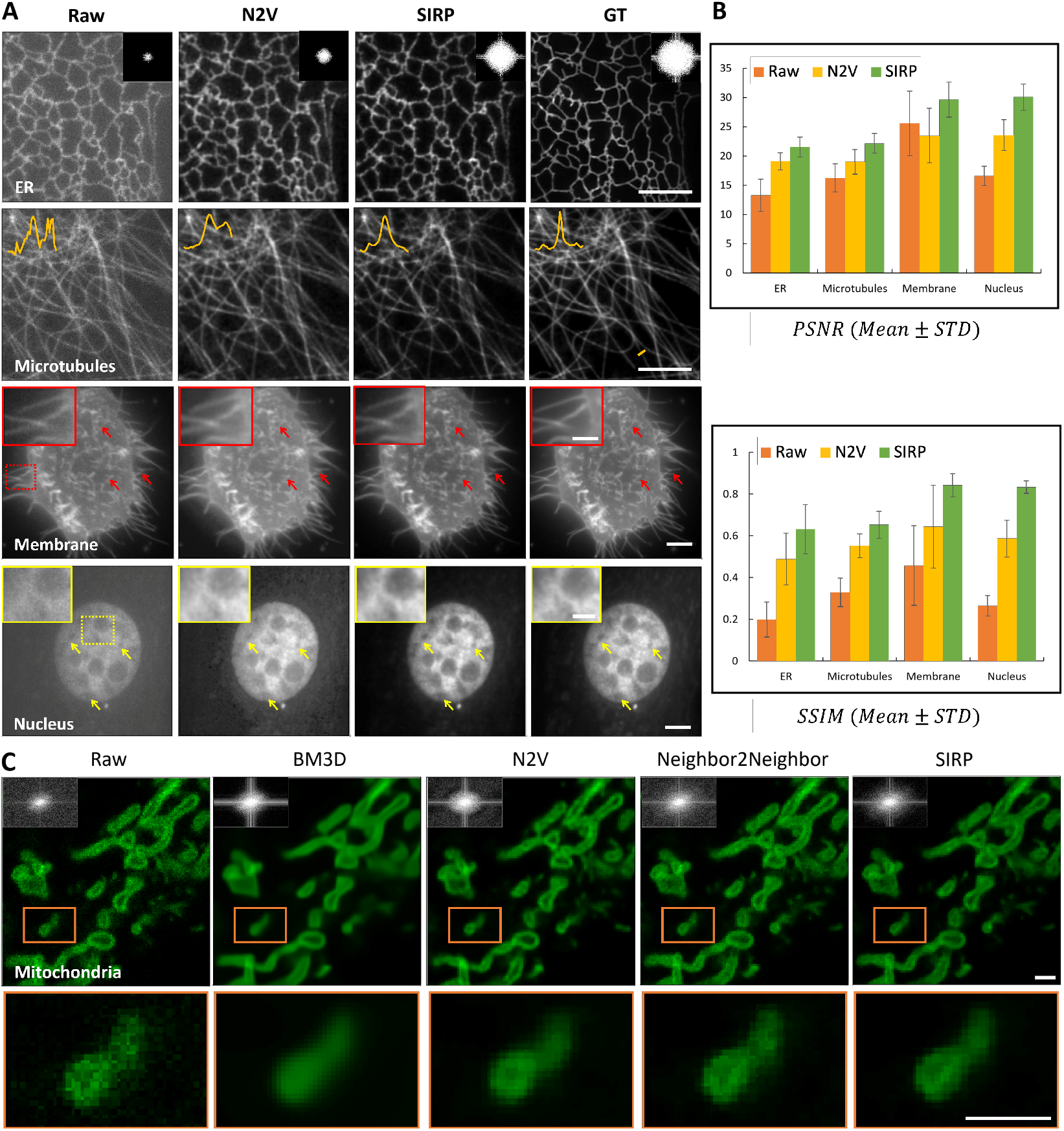
Denoising comparison between SIRP with different methods using different biological samples. (A) Comparison between SIRP and Noise2Void (N2V) on four real-acquired biological datasets. From top to bottom, the datasets are ER, Microtubules, Membrane, and Nucleus. For the ER dataset, Fourier spectra of selected regions are displayed. For the Microtubules dataset, line profiles of selected regions are shown. The magnified views and arrows indicate regions where SIRP outperforms N2V. Scale bars: 5 *µ*m (ER, Microtubules); 2 *µ*m (Membrane, Nucleus). (B) Quantitative evaluation of PSNR and SSIM across the four datasets with respect to their ground truths (±mean standard deviation, tested over 20 images each group). (C) Comparison among BM3D, N2V, Neighbor2Neighbor, and SIRP on mitochondrial images. The Fourier spectra of each restored image are shown in the upper-left corner, and magnified views of ROIs are displayed below. Scale bars: 1 *µ*m.

The Membrane and Nucleus datasets were acquired using an IX83 inverted fluorescence microscope [30], low-SNR (Raw) and high-SNR(GT) images were obtained by varying the exposure time. The magnified views and arrows highlight regions where SIRP maintains intracellular shapes and boundary integrity that are partially lost or still blurred in Noise2Void results. For nucleus images, SIRP provided a cleaner background and clearer nuclear pores compared with N2V results.

SIRP consistently outperforms Noise2Void in both denoising quality and structural fidelity. Overall, SIRP produces cleaner backgrounds, clearer structural boundaries, and higher contrast reconstructions. These advantages are reflected in both qualitative observations and quantitative metrics.

These qualitative differences align with the quantitative evaluation in Fig. 4B. Across all datasets, SIRP achieves higher PSNR and SSIM scores with smaller error bars, demonstrating superior reconstruction accuracy and robustness over N2V.

We additionally compared SIRP with several representative denoising approaches on mitochondrial images acquired using a confocal microscope [22] (× 100 oil-immersion objective, NA 1.45), as shown in Fig. 4C. This dataset does not have the no-noise reference. We mainly do qualitative analysis. While all methods reduce noise to some extent, their performances varied. Magnified views reveal that BM3D tended to oversmooth fine details, and Neighbor2Neighbor caused structural disturbance of the mitochondrial membrane. Although Noise2Void (N2V) also achieved good denoising performance, its restored frequency spectrum exhibits a discrepancy in the orientation of its spectral distribution compared to the raw image, suggesting potential artifacts. Compared with others, SIRP retained more accurate structural details and preserves high-frequency information more effectively.

### 3.4 Key Factors Affecting SIRP Performance

To comprehensively analyze the factors influencing reconstruction quality, we performed ablation experiments on two key components of the Image Representation Module, contrast enhancement and the activation/encoding strategy used for prior generation.

The contrast level of the prior plays an essential role in shaping both the background fidelity and the overall reconstruction quality. In our ablation study, the low-contrast (LC) prior corresponded to a *γ* of 1.0, which means no contrast enhancement, whereas the high-contrast (HC) prior was obtained using a *γ* of 1.5. Priors with low contrast tend to produce brighter backgrounds and more blurry structural boundaries, whereas enhancing the contrast helps produce cleaner backgrounds that better match the ground truth (Fig. 5A). Consistently, both PSNR and SSIM increase under the enhanced-contrast setting, highlighting the effect of contrast enhancement and the sensitivity of the reconstruction results to the quality of the image representation prior.

**Fig. 5.**
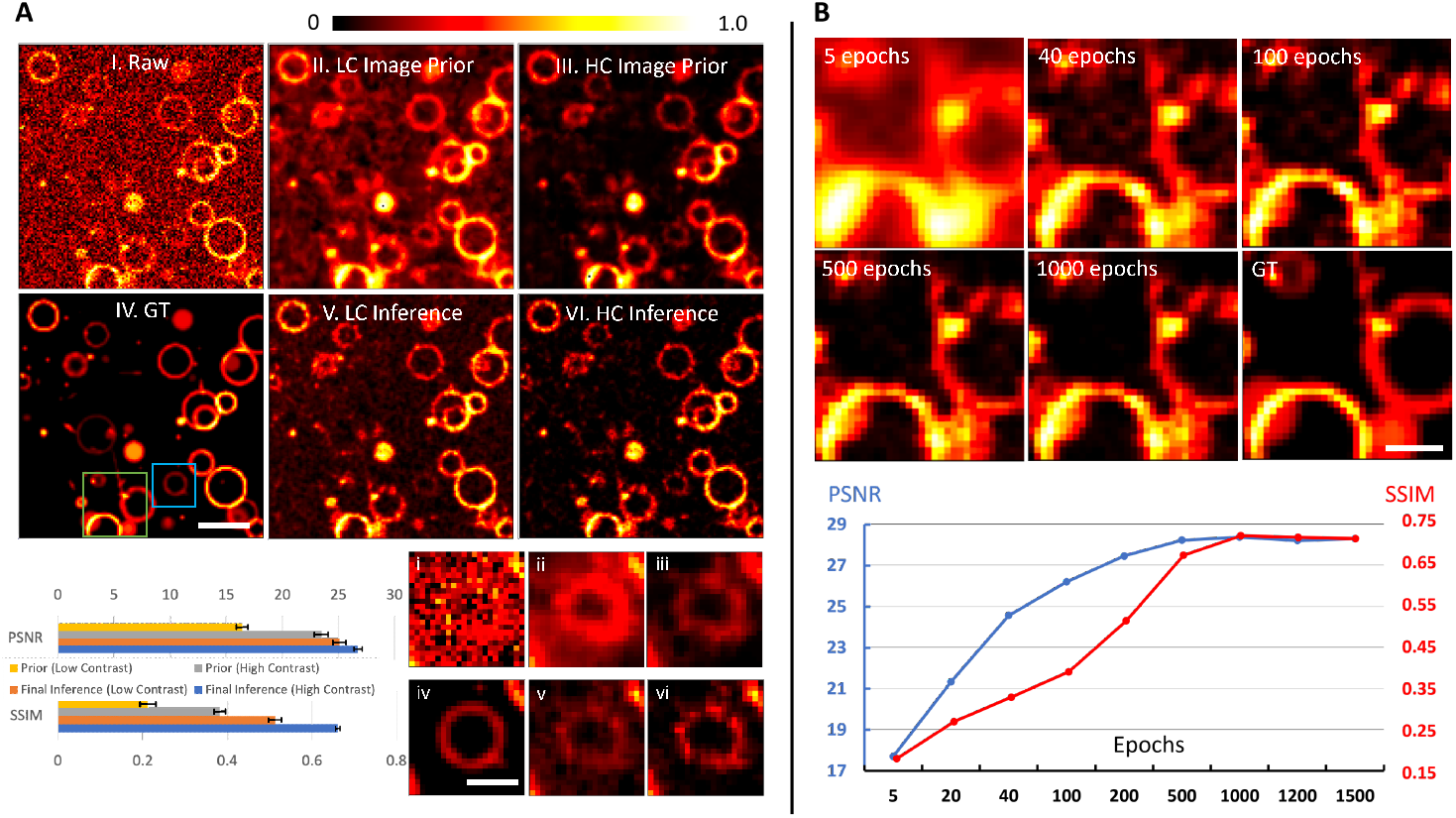
Effect of contrast enhancement in the Image Representation Module and convergence behavior of SIRP, experimented on synthetic images with SNR∼ 10dB. (A) Contrast-enhancement component enables better noise suppression. With contrast enhancement enabled, the image prior exhibits high contrast (HC), whereas disabling it yields a low-contrast (LC) prior. Quantitative metrics (PSNR and SSIM) for each setting are reported accordingly. Insets at the bottom-right show the corresponding ROIs, corresponding to the blue boxed regions in the full images. The pseudo-color normalized intensity color bar (top) is shared by both (A) and (B). Scale bars: 25 pixels in original-scale image; 6 pixels in magnified views. (B) Convergence analysis with respect to the training epoch number of the U-Net. Displayed region corresponds to the green boxed area in (A). The denoising effect gradually improves as the iterations proceed. The plots below show how PSNR and SSIM evolve as the training progresses, illustrating the convergence of the denoising network. Scale bar: 15 pixels.

The choices for encoding and activation also determine the representational capacity of the INR and therefore influence the final reconstruction quality. Consequently, identifying the most effective combination of these structural elements is essential. To investigate the most effective combination of element selection, we systematically evaluated the performance of different coordinate encoding strategies, including raw coordinates, Fourier Reparameterization (FR), and Positional Encoding (PE), in conjunction with distinct activation functions (ReLU and Sine). The Sine-based models tested here follow the SIREN formulation [26], where the INR is constructed using raw input coordinates together with sinusoidal activations. Although SIREN-based architecture often achieves superior performance in conventional INR tasks due to its strong ability to represent high-frequency signals, it tends to overfit the stochastic noise present in low-SNR images, thereby reducing the reconstruction quality. In contrast, ReLU activations, with their piecewise-linear nature, inherently impose a smoother functional prior. When paired with appropriately scaled positional encoding, the network captures essential structural details while suppressing high-frequency noise. This balance between structural fidelity and noise filtering explains the superior performance of the ReLU + Positional Encoding configuration. As detailed in Table 1, the configuration employing ReLU with Positional Encoding yielded the highest quantitative metrics on simulated images at an SNR of 10 dB.

**Table 1.** Impact of different encoding schemes and activation functions for generating the prior on the final inference.

| | PSNR (Mean±STD) | | SSIM (Mean±STD) | |
| --- | --- | --- | --- | --- |
|  | Prior | Final Inference | Prior | Final Inference |
| ReLU | 26.31±1.37 | 28.38±1.32 | 0.55±0.01 | 0.71±0.01 |
| ReLU+FR | 22.22±1.21 | 28.21±1.28 | 0.19±0.02 | 0.67±0.02 |
| **ReLU+PE** | **26.57±1.03** | **28.47±0.98** | **0.61±0.01** | **0.73±0.01** |
| Sine | 24.88±0.89 | 28.05±1.02 | 0.39±0.01 | 0.61±0.01 |
| Sine+FR | 24.33±1.15 | 28.29±1.19 | 0.36±0.01 | 0.64±0.01 |

**Note.** FR = Fourier Reparameterization; PE = Positional Encoding.

Another important factor is the total number of U-net training iterations. We further analyzed the convergence behavior of SIRP. As training proceeded, backgrounds of reconstruction gradually became cleaner, dark regions went toward their true intensity, and edge-related artifacts were reduced until convergence (Fig. 5B). The accompanying PSNR and SSIM curves similarly reach plateaus after 1000 epochs. Therefore, for most experiments, we set the training number as 1000 epochs.

## 4. Discussion

In this work, we proposed a novel self-supervised framework that effectively integrates image representation prior with a neighbor-sampling strategy for image denoising. By leveraging INR’s continuous image representation capability, our method generates smooth, structurally coherent pre-denoised priors that guide the end-to-end network training, overcoming the limitation of requiring clean ground-truth images. This makes our approach particularly suitable for challenging low-SNR microscopy scenarios where acquiring clean data is impractical. Extensive experiments on both synthetic images and real-acquired fluorescence microscopy data demonstrate that our method achieves superior denoising performance compared to both traditional denoising methods and baseline self-supervised approaches, producing clearer structural details, cleaner backgrounds, and significantly reduced noise-induced artifacts.

While integrating INRs offers notable advantages in continuous signal representation and structural smoothness, it also introduces several practical limitations. A key limitation of INR-based priors lies in the per-image training requirement: although this step is performed only during prior construction and does not affect inference speed, it nevertheless increases the overall training time when processing large datasets. Beyond computational considerations, INRs also exhibit an inherent trade-off between denoising strength and structural fidelity, especially under extremely low SNR conditions. When noise dominates the measurements, the INR may oversmooth fine details or fail to fully recover high-frequency structures. Our end-to-end framework is designed to mitigate these limitations, but the balance between prior smoothness and information preservation remains an important consideration.

Despite these challenges, combining INR priors with self-supervised training provides a powerful approach for fluorescence image denoising. Future work will focus on extending the framework to 3D volumetric and 4D spatiotemporal datasets, and leveraging shared representations across spatial and temporal dimensions to enhance scalability and robustness, optimizing INR architectures with hybrid designs. Overall, SIRP provides a robust solution to overcome the noise degradation in fluorescent microscopy, and the image representation prior strategy can be extended to other self-supervised frameworks, broadening the practical applicability of image restoration algorithms.

## Notes

https://gigadb.org/dataset/100888

https://figshare.com/articles/dataset/BioSR/13264793

https://www.nature.com/articles/s41592-023-02058-9

### Competing Interest Statement

The authors have declared no competing interest.

### Funding

- National Natural Science Foundation of China: 62427807, 62475232, 62405269, 6240030418
- Talent Program of Zhejiang Province: 2021R51004
- Leading Innovative and Entrepreneur Team Introduction Program of Zhejiang: 2024R01001

## References

1. J. W. Lichtman and J.-A. Conchello, “Fluorescence microscopy,” Nat. Methods 2, 910–919 (2005).

2. A. Rizk, G. Paul, P. Incardona, et al., “Segmentation and quantification of subcellular structures in fluorescence microscopy images using squassh,” Nat. Protoc. 9, 586–596 (2014).

3. S. Wäldchen, J. Lehmann, T. Klein, et al., “Light-induced cell damage in live-cell super-resolution microscopy,” Sci. Reports 5, 15348 (2015).

4. “Phototoxicity revisited,” Nat. Methods 15, 751 (2018).

5. J. Icha, M. Weber, J. C. Waters, and C. Norden, “Phototoxicity in live fluorescence microscopy, and how to avoid it,” BioEssays 39 (2017).

6. R. A. Hoebe, C. J. van Noorden, and E. M. Manders, “Noise effects and filtering in controlled light exposure microscopy,” J. Microsc. 240, 197–206 (2010).

7. R. A. Hoebe, H. T. Van der Voort, J. Stap, et al., “Quantitative determination of the reduction of phototoxicity and photobleaching by controlled light exposure microscopy,” J. Microsc. 231, 9–20 (2008).

8. Y. Wu and H. Shroff, “Multiscale fluorescence imaging of living samples,” Histochem. Cell Biol. 158, 301–323 (2022).

9. M. Mora-González, J. Muñoz-Maciel, F. J. Casillas, et al., “Image processing for optical metrology,” MATLAB–a ubiquitous tool for practical engineer. 1st ed. Rijeka, Croat. InTech pp. 523–546 (2011).

10. C. Tomasi and R. Manduchi, “Bilateral filtering for gray and color images,” in Sixth international conference on computer vision (IEEE Cat. No. 98CH36271), (IEEE, 1998), pp. 839–846.

11. A. Buades, B. Coll, and J.-M. Morel, “Non-local means denoising,” Image processing on line 1, 208–212 (2011).

12. L. Yang, R. Parton, G. Ball, et al., “An adaptive non-local means filter for denoising live-cell images and improving particle detection,” J. Struct. Biol. 172, 233–243 (2010).

13. S. Zheng, M. Koyama, and J. Mertz, “Multiplane hilo microscopy with speckle illumination and non-local means denoising,” J. biomedical optics 28, 116502 (2023).

14. C.-W. Chang and M.-A. Mycek, “Total variation versus wavelet-based methods for image denoising in fluorescence lifetime imaging microscopy,” J. biophotonics 5, 449–457 (2012).

15. K. Zhang, W. Zuo, Y. Chen, et al., “Beyond a gaussian denoiser: Residual learning of deep cnn for image denoising,” IEEE transactions on image processing 26, 3142–3155 (2017).

16. O. Ronneberger, P. Fischer, and T. Brox, “U-net: Convolutional networks for biomedical image segmentation,” in International Conference on Medical image computing and computer-assisted intervention, (Springer, 2015), pp. 234–241.

17. A. Krull, T.-O. Buchholz, and F. Jug, “Noise2void-learning denoising from single noisy images,” in Proceedings of the IEEE/CVF conference on computer vision and pattern recognition, (2019), pp. 2129–2137.

18. J. Lehtinen, J. Munkberg, J. Hasselgren, et al., “Noise2noise: Learning image restoration without clean data,” arXiv preprint arXiv:1803.04189 (2018).

19. T. Huang, S. Li, X. Jia, et al., “Neighbor2neighbor: Self-supervised denoising from single noisy images,” in Proceedings of the IEEE/CVF conference on computer vision and pattern recognition, (2021), pp. 14781–14790.

20. X. Tian, Q. Wu, H. Wei, and Y. Zhang, “Noise2sr: Learning to denoise from super-resolved single noisy fluorescence image,” (2022).

21. J. Batson and L. Royer, “Noise2self: Blind denoising by self-supervision,” in International conference on machine learning, (PMLR, 2019), pp. 524–533.

22. G. Zhang, X. Li, Y. Zhang, et al., “Bio-friendly long-term subcellular dynamic recording by self-supervised image enhancement microscopy,” Nat. Methods 20, 1957–1970 (2023).

23. S. Laine, T. Karras, J. Lehtinen, and T. Aila, “High-quality self-supervised deep image denoising,” Adv. neural information processing systems 32 (2019).

24. D. Ulyanov, A. Vedaldi, and V. Lempitsky, “Deep image prior,” in Proceedings of the IEEE conference on computer vision and pattern recognition, (2018), pp. 9446–9454.

25. C. Zhang and K. S. Yen, “A survey on deep image prior for image denoising,” Digit. Signal Process. 163, 105235 (2025).

26. V. Sitzmann, J. N. P. Martel, A. W. Bergman, et al., “Implicit neural representations with periodic activation functions,” (2020).

27. B. Mildenhall, P. P. Srinivasan, M. Tancik, et al., “Nerf: Representing scenes as neural radiance fields for view synthesis,” Commun. ACM 65, 99–106 (2021).

28. H. Zou and T. Hastie, “Regularization and variable selection via the elastic net,” J. Royal Stat. Soc. Ser. B: Stat. Methodol. 67, 301–320 (2005).

29. C. Qiao and D. Li, “BioSR: a biological image dataset for super-resolution microscopy,” (2020).

30. G. M. Hagen, J. Bendesky, R. Machado, et al., “Fluorescence microscopy datasets for training deep neural networks,” GigaScience 10, giab032 (2021).

